# Lipid nanoparticle structure and delivery route during pregnancy dictates mRNA potency, immunogenicity, and health in the mother and offspring

**DOI:** 10.1101/2023.02.15.528720

**Authors:** Namit Chaudhary, Alexandra N. Newby, Mariah L. Arral, Saigopalakrishna S. Yerneni, Samuel T. LoPresti, Rose Doerfler, Daria M. Strelkova Petersen, Bethany Fox, Tiffany Coon, Angela Malaney, Yoel Sadovsky, Kathryn A. Whitehead

**Author notes:** Corresponding author: Kathryn A. Whitehead.

## Abstract

Treating pregnancy-related disorders is exceptionally challenging because many small molecule drugs on the market may cause maternal and fetal toxicity. This potential danger has hindered the development and clinical evaluation of new drugs for several decades. Lipid nanoparticle (LNP)-based RNA therapies with high delivery efficacy, favorable immune response, and minimal transplacental transport can quell maternal-fetal toxicity concerns and propel the development of pregnancy-safe drugs. To this extent, we report potent LNP structures that robustly deliver mRNA to maternal organs and placenta. Using structure-function analysis, we show that LNP efficacy is influenced by the polyamine headgroup, and toxicity is governed by the acrylate tail. Our lead nanoparticle shows robust protein expression via multiple clinically relevant administration routes in pregnant mice. In the placenta, it transfects trophoblasts, endothelial cells, and immune cells. Further, by varying ionizable lipid structure, we demonstrate that LNP immunogenicity affects organ expression and pup health during pregnancy. Immunogenic LNPs show lower efficacy in lymphoid organs in an IL-1β dependent manner in pregnant mice. Further, pro-inflammatory immune responses provoke the infiltration of adaptive immune cells in the placenta and restrict pup growth after birth. Together, our results provide a mechanistic basis for designing safe and potent LNPs that can be administered during pregnancy.

## Introduction

Physiological and immunological changes during pregnancy can exacerbate pre-existing maternal health conditions, and induce emergent conditions, thus increasing the risk of maternal medical and obstetrical complications (1). The impact of resultant adverse events on maternal and fetal health can be devastating. For example, the hypertensive disorder, preeclampsia, affects 5–10% pregnancies and causes over 75,000 maternal and 500,000 infant deaths globally each year (2). Additionally, pregnant people with SARS-CoV-2 infection are twice as likely to need intensive care, invasive ventilation, and extracorporeal membrane oxygenation compared to non-pregnant people (3).

Although hypertension, viral infections, and other conditions are often manageable in non-pregnant people, they can be life-threatening during pregnancy because many relevant drugs have not been proven safe. Safety data are scarce because pregnant people have been commonly excluded from clinical trials (4). As of 2022, some international clinical trial registries report that pregnant people were included in a mere 0.15% of registered clinical trials (5). Because of lack of safety data, many commonly used drugs, including the analgesic ibuprofen, the antidepressants clonazepam and lorazepam, the antimalarial primaquine, and the antibiotics ciprofloxacin, levofloxacin, and trimethoprim are restricted during pregnancy (6). Additionally, several drugs, such as the blood thinner warfarin, and the antimalarial primaquine are restricted because of known toxicity concerns (6). These small molecule drugs rapidly transport across the placental barrier due to their small size, and may impair fetal development.

To prevent fetal accumulation, lipid nanoparticle (LNP) formulations may be safer drug delivery alternatives because the large nanoparticle coating (~100 nm) hinders transplacental transport (7). Because they are degradable and highly biocompatible, LNPs typically clear from the bloodstream rapidly without long-term accumulation in organs (8, 9). Additionally, the encapsulated RNA therapeutics are advantageous over small molecules because they target dysregulated proteins at the genetic level with higher specificity (10). Indeed, clinical trial data suggest that the mRNA-LNP-based SARS-CoV-2 from Moderna and Pfizer–BioNTech have excellent safety profiles in pregnant people (11). Compared to unvaccinated individuals, mRNA vaccines reduced the risk of severe COVID-19 illness in pregnant individuals (12), and the risk of COVID-19 hospital admission among their infants younger than six months of age (13). Moreover, vaccination did not increase the risk of pregnancy-related complications, including maternal death, stillbirth, preterm birth, placental abruption, and postpartum hemorrhage (14).

These results are promising; however, critical design parameters that will extend the success of mRNA vaccines to other therapeutic LNP platforms in pregnant people are scarce. Specifically, there is a lack of understanding of changes in LNP efficacy, placental transport, and immune response during pregnancy. To overcome this design challenge, we report LNP structures that potently deliver mRNA to the placenta and non-reproductive maternal organs with no fetal delivery in mice. LNPs transfect several major placental cell types, including trophoblasts, endothelial cells, and placental immune cells. We also show that immunostimulation alters LNP organ tropism and inhibits pup development. Together, these experiments identify potent LNP structures and design criteria for potent and safe mRNA delivery during pregnancy in a murine model.

## Results

### LNP headgroup and tailgroup govern mRNA delivery and toxicity, respectively

To design LNPs that deliver mRNA to the placenta, we first assessed the delivery efficacy of a library of LNPs in BeWo b30 cells. BeWo b30 are human choriocarcinoma cells. They express several critical differentiation markers (15) that model the placental trophoblast barrier that separates maternal and fetal blood circulation. They are a good choice to study the uptake (16), transport (17), and efflux (18) of drugs across the placental trophoblasts *in vitro*.

Our library contained 260 LNPs formulated using ionizable lipids that we synthesized combinatorically from 20 amine heads and 13 acrylate tails (Fig. S1). To test efficacy, we encapsulated mRNA encoding firefly luciferase in LNPs, and added them to BeWo b30 cells. The predominant determinant of mRNA delivery potency was the headgroup of the ionizable lipid, with amines 306, 500, 514, and 516 producing the highest levels of protein expression (Fig. 1A).

**Figure 1:**
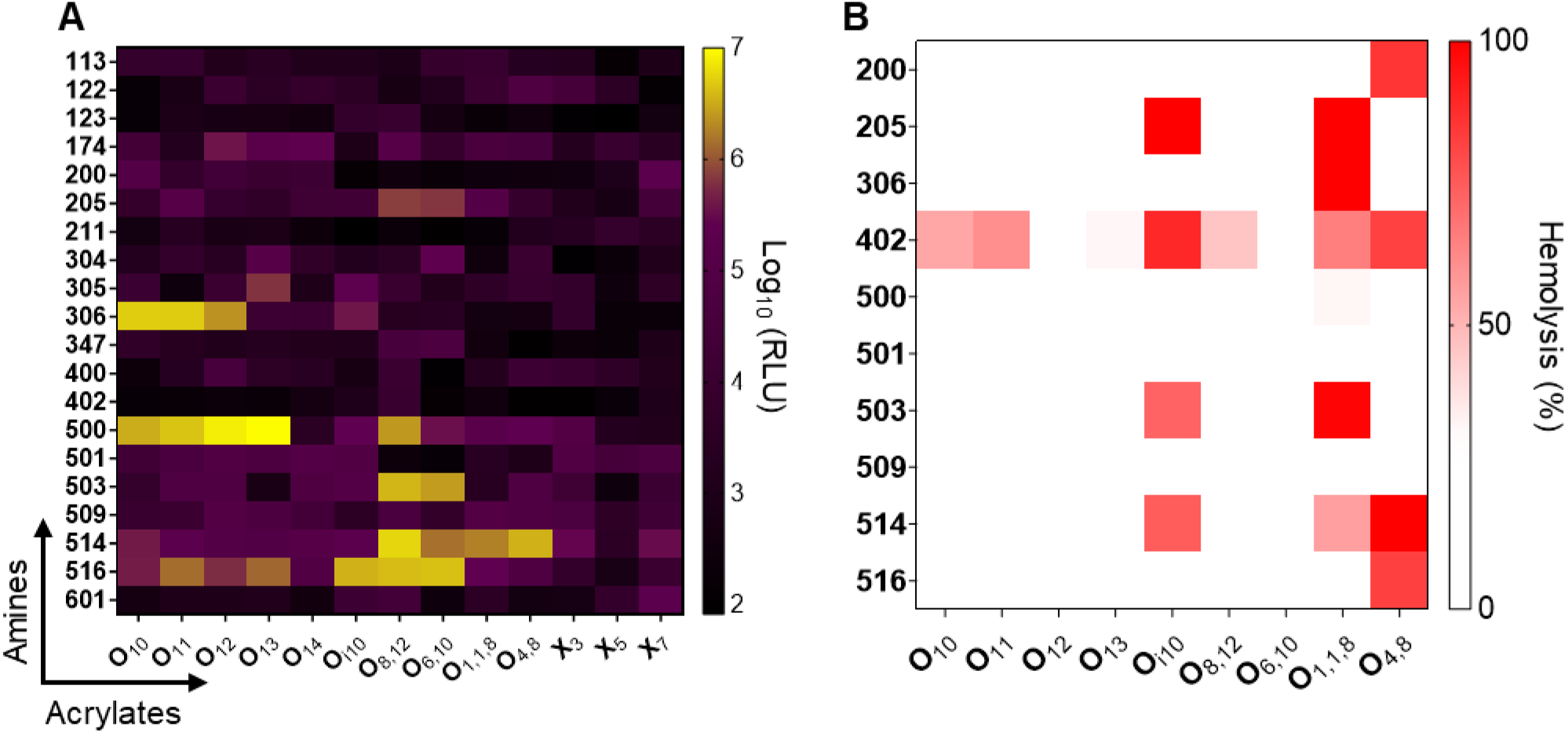
Ionizable lipid structure influences mRNA delivery in human placental trophoblasts *in vitro*. (A) 260 ionizable lipids generated from 20 amine headgroups (y-axis) and 13 alkyl acrylate tails (x-axis) were formulated into lipid nanoparticles encapsulating firefly luciferase mRNA. Nanoparticles were incubated with BeWo b30 cells for 24 hours at a dose of 100 ng mRNA, and luminescence intensity was measured in relative light units (RLU), with brighter colors indicating better transfection. (B) Nanoparticle-induced hemolysis of human red blood cells was measured as a proxy for toxicity in the bloodstream. Red Blood cells were incubated with blank LNPs at an equivalent mRNA dose of 2500 ng for 1.5 hours, and hemolysis was assessed by measuring absorbance in the supernatant. In both panels, the heat map colors represent median values (n = 4).

Because safety is equally important as efficacy for the successful clinical translation, we next assessed toxicity with a lysis assay to rule out LNPs that destroy red blood cells. In this assay, toxic LNPs would burst red blood cells at physiological pH and release the chromogenic heme complex into the supernatant. LNP hemolytic activity was quantified by measuring the color intensity of the supernatant using a spectrophotometer. We incubated 90 LNP structures with red blood cells and observed that the majority of the LNPs did not lyse the cells. However, some LNP structures were hemolytic, including those containing the 402 amine head group (Fig. 1B). Interestingly, some LNPs containing branched tails (e.g., O_i10_ and O_1,1,8_) were more likely to induce hemolysis than their linear tail counterparts O_10_ and O_11_ (Fig. 1B). To minimize safety concerns, we eliminated hemolytic LNPs from *in vivo* studies.

### LNPs don’t penetrate the trophoblast barrier *in vitro*

Next, we gauged if viable LNPs penetrate the trophoblast barrier. For this, we cultured BeWo b30 cells on semipermeable Transwell™ inserts for a week so that they fuse together and form a “syncytialized” trophoblast barrier *in vitro* (19, 20). Barrier formation was assessed by measuring transepithelial electrical resistance (TEER) across the apical and basolateral compartments of the insert, which increases with barrier function. In addition to the b30 clones, we used non-fusogenic parental BeWo cells as a negative control (20). As expected, TEER increased only for syncytium-forming BeWo b30 cells (Fig. 2A), with TEER exceeding 100 and 200 Ohm-cm^2^ after five and seven days, respectively. TEER >100 Ohm-cm^2^ suggests successful barrier formation in BeWo b30 cells (17).

**Figure 2:**
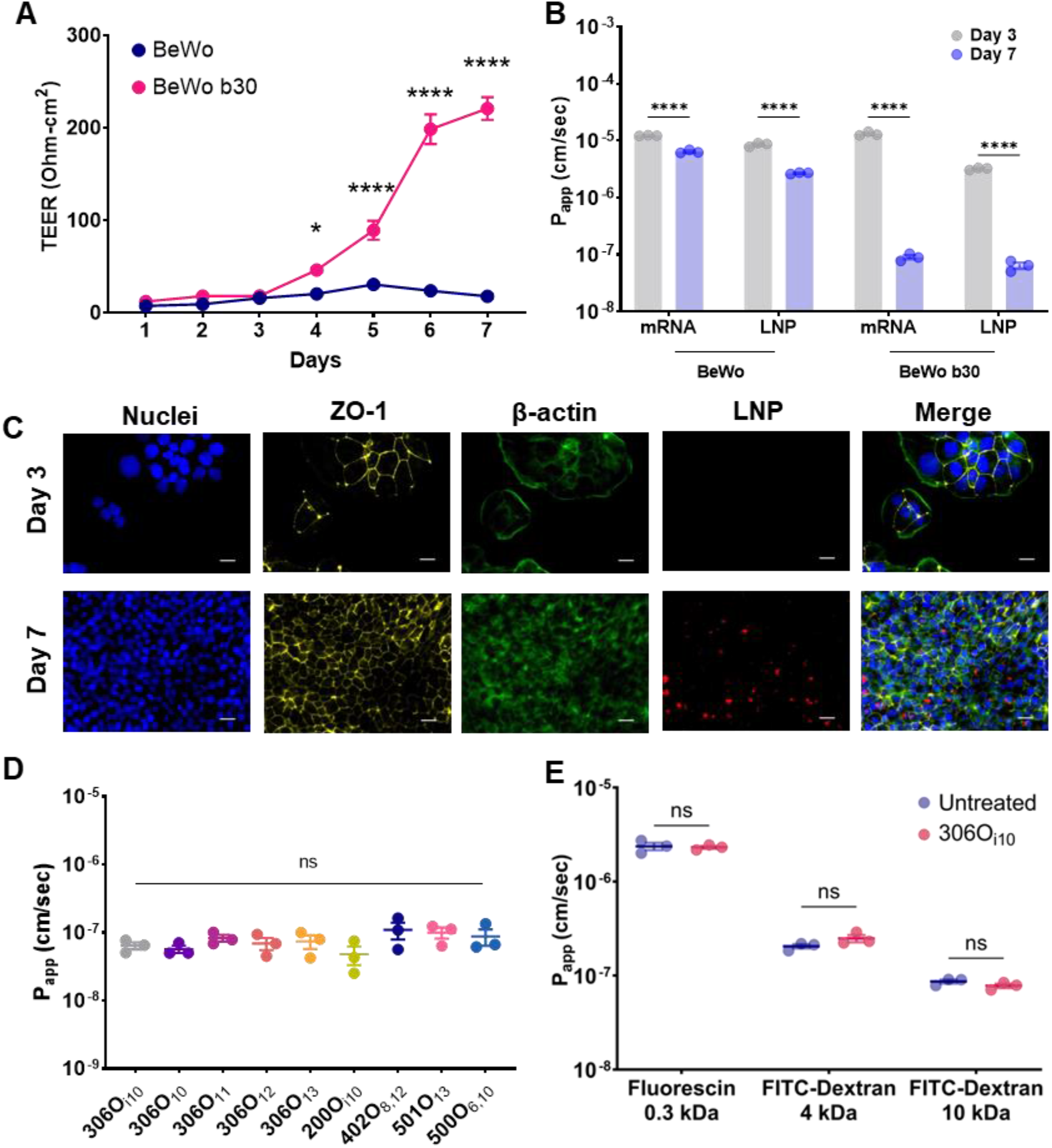
The placental barrier inhibits lipid nanoparticle transport *in vitro*. (A) When grown on Transwell plates, human trophoblast BeWo b30 clone cells formed a representative trophoblast barrier, whereas parent BeWo cells did not. Monolayer formation was marked by increases transepithelial electrical resistance (TEER, n = 12). (B) The apparent permeability (P_app_) of naked and LNP-encapsulated Cy5-labeled mRNA was assessed as a function of time across BeWo and BeWo b30 monolayers. Fully formed BeWo b30 monolayers inhibited naked and encapsulated mRNA transport (n = 3) (C) Nanoparticle uptake by BeWo b30 monolayers was imaged using immunofluorescence microscopy: DAPI (blue); anti-ZO-1 antibody (yellow); Phalloidin (green); Cy5-tagged mRNA (red). All scale bars are 20 μm. (D) On day 7 of BeWo b30 monolayer development, the apparent permeability of LNPs was not affected by ionizable lipid chemistry. (n = 3) (E) The apparent permeability of 0.3, 4, and 10 kDa fluorescent marker molecules was assessed in the presence or absence of 306O_i10_ LNPs, which showed that LNPs did not act as permeation enhancers (n = 3). Error bars represent s.e.m. ns, *, and **** represent non-significant, p < 0.05, and p < 0.0001, respectively. Panels A and D were analyzed using one-way ANOVA with Dunnett’s and Tukey’s post-hoc analysis, respectively, and panel C and E used two-way ANOVA with Tukey’s and Sidak’s post-hoc analysis, respectively.

To test barrier function, we quantified transport kinetics by measuring the apparent permeability (P_app_) of Cy5-labelled naked mRNA or mRNA-LNPs in BeWo and BeWo b30 cells. For this, we chose 306O_i10_ LNPs because they are not hemolytic (Fig. 1B) and have been previously shown to be exceptionally potent *in vivo* (21, 22). We observed that parental BeWo cells reduced P_app_ by ~2-fold for naked mRNA and ~5-fold for 306O_i10_ LNPs between days 3 and 7 (Fig. 2B). In comparison, syncytialized BeWo b30 cells reduced mRNA and LNP transport by ~136-fold and ~192-fold, respectively (Fig. 2B).

Fluorescence microscopy revealed that BeWo b30 cells developed a confluent cell network by day 7. Compared to day 3, the monolayers robustly expressed the barrier-forming tight junction protein zonula occludens-1 (23) and took up LNPs to a greater degree (Fig. 2C). This suggests that the syncytialized BeWo b30 “catch” LNPs, preventing them from diffusing between the cells. Further, we observed that BeWo b30 cells inhibited transport regardless of LNP structure, with a total of nine LNPs showing no statistical differences in transport rate (Fig. 2D).

In addition to restricting fetal exposure to pathogen and foreign substances, the trophoblasts also facilitates the exchange of molecules of varied size, ranging from small-molecule nutrients (24), such as glucose, fatty acids, and amino acids to macromolecular peptides and proteins (25). For healthy fetal development, it is critical that LNPs don’t hinder the exchange of such molecules. To show this, we exposed BeWo b30 cells to differently sized marker molecules, including 300 Da fluorescein and 4 kDa and 10 kDa FITC-Dextrans. We observed that 306O_i10_ did not alter transport kinetics of any marker molecule (Fig. 2E). Together, these data show that LNPs don’t penetrate the trophoblast barrier or hinder transport of other small- or macro-molecules *in vitro*.

### Immune response shifts LNP organ tropism in pregnant mice

Having shown that LNPs can safely deliver mRNA to placental cells *in vitro,* we asked if LNPs recapitulate this action *in vivo*. For this, we formulated 306O_i10_ LNPs encapsulating firefly luciferase mRNA and injected them intravenously (IV) in CD-1 mice on day 14 of pregnancy and in age-matched non-pregnant females Mice gestate on average for 19-21 days, and the placenta is functional by day 12.5 (26, 27). Therefore, day 14 corresponds to the early phases of second trimester in a human pregnancy (28). We observed dose-dependent LNP delivery to the placenta (Fig. 3A, 3B), with no observable signal in the fetus (Fig. 3C). Among non-reproductive organs, firefly luciferase was expressed to the greatest extent in the liver and spleen (Fig. 3D, 3E). Additionally, there were no shifts in non-reproductive organ tropism between non-pregnant and pregnant mice at any dose (Fig. 3D, 3E).

**Figure 3:**
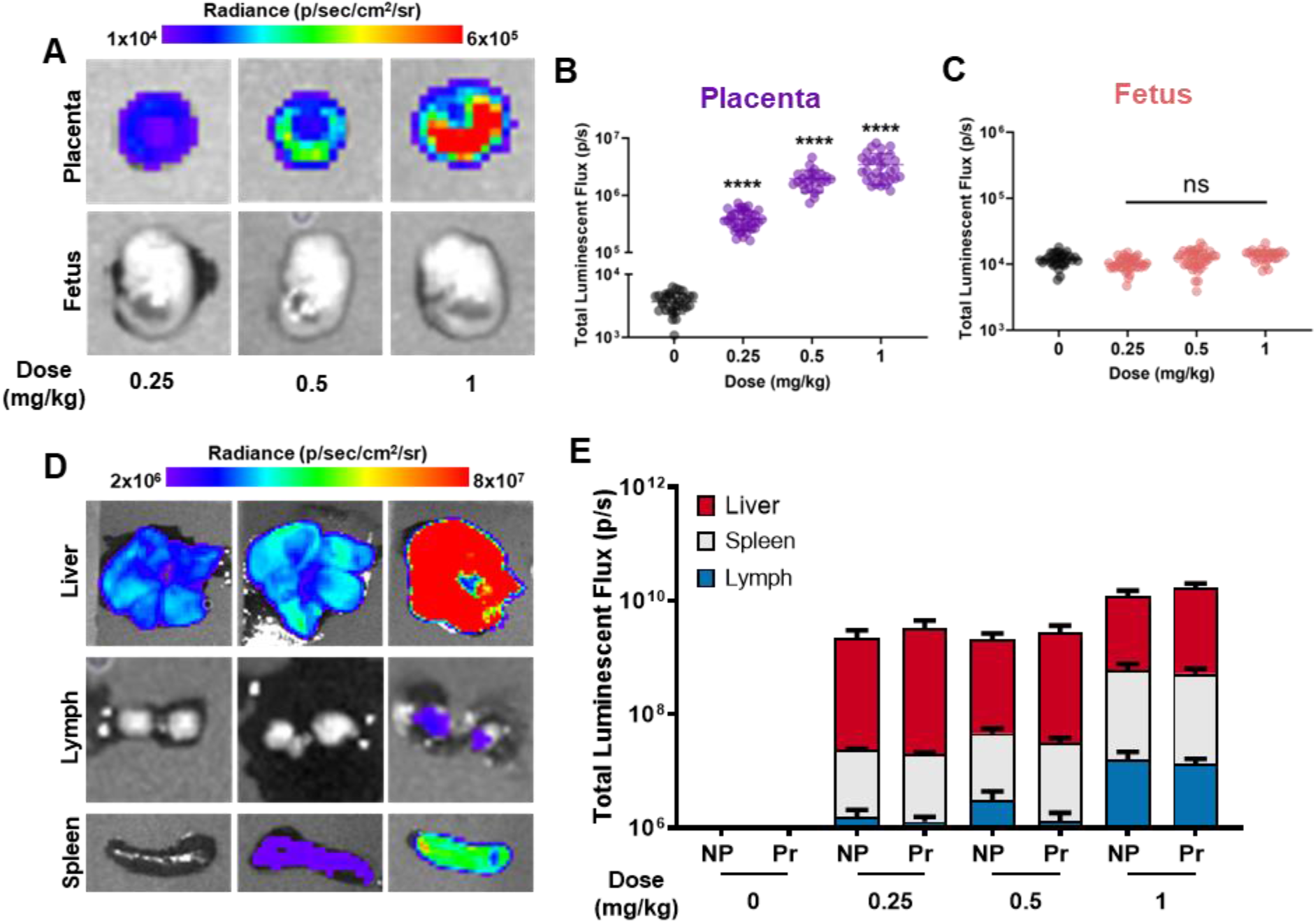
306O_i10_ LNPs elicit dose-dependent mRNA delivery to the placenta. 306O_i10_ LNPs encapsulating luciferase mRNA were injected intravenously in non-pregnant (NP) or pregnant (Pr) CD-1 mice. Four hours later, mice were euthanized, and total luminescent flux was measured in (A-C) placenta and fetus and (D-E) non-reproductive organs using IVIS. Error bars represent s.e.m. with n = 3 dams per treatment group and **** representing p < 0.0001, according to one-way ANOVA with Dunnett’s post-hoc analysis.

In addition to IV delivery, we considered other clinically relevant delivery routes, including intramuscular (IM) and intraperitoneal (IP) administration. Intramuscular injections are the preferred route for prophylactic RNA vaccines (29, 30), including the SARS-CoV-2 vaccines (31, 32), while intraperitoneal injections are commonly used clinically to deliver therapeutics to intraabdominal organs (33–35). We saw robust mRNA delivery for all administration routes, with IV delivery eliciting the highest placental delivery, followed by IP and IM (Fig. 4A, 4B). None of the administration routes resulted in luminescence in the fetal compartment (Fig. 4A, 4C).

**Figure 4:**
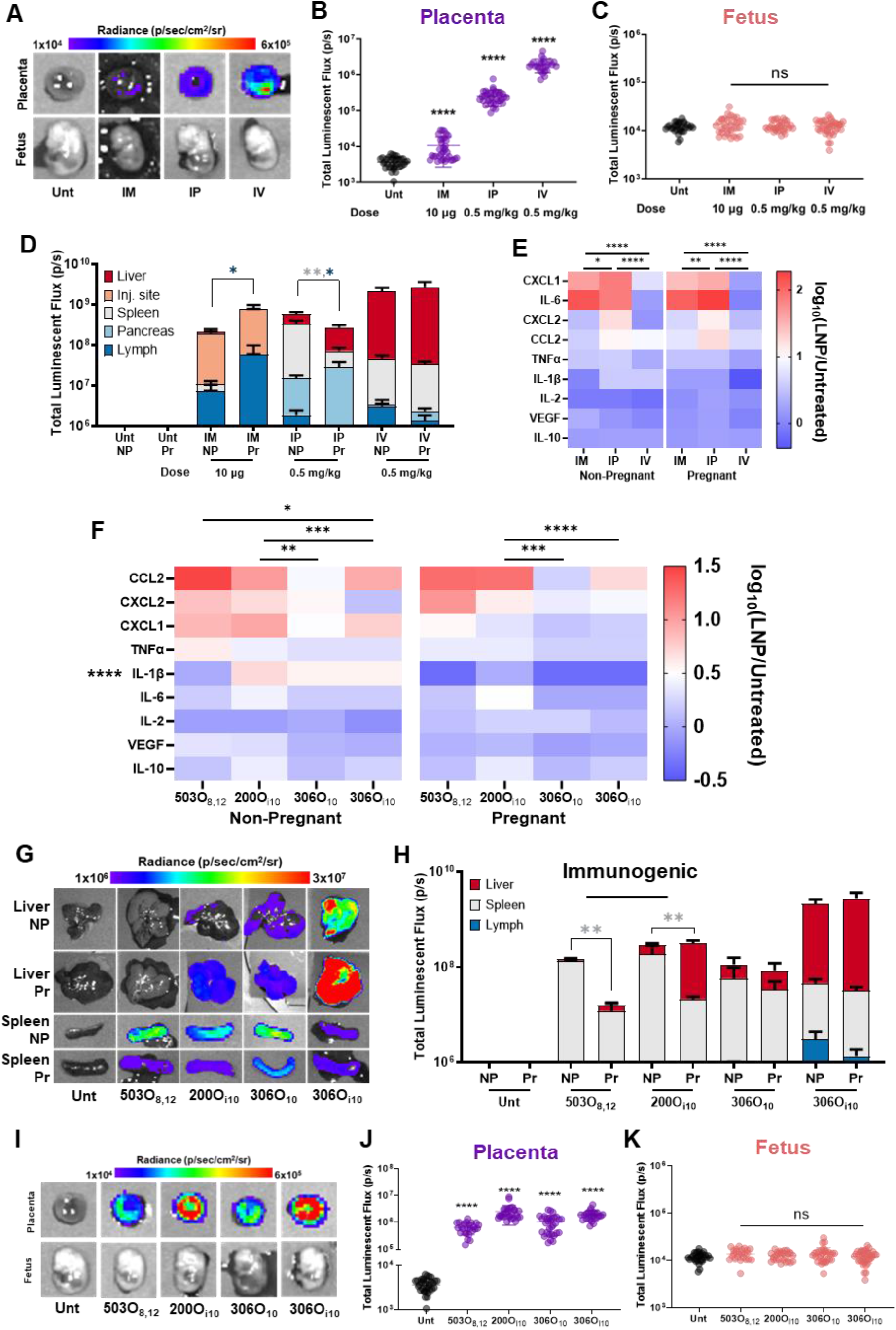
Lipid nanoparticles deliver mRNA to the placenta without delivery to the fetus. LNPs encapsulating firefly luciferase mRNA were injected intramuscularly (10 μg), intraperitoneally (0.5 mg/kg), or intravenously (0.5 mg/kg) in non-pregnant or pregnant CD-1 mice on the 14^th^ day of pregnancy. Four hours later, mice were euthanized, and their organs were imaged for luminescence using IVIS. (A-C) Efficacy using lipid 306O_i10_ was quantified in the placenta and fetus for pregnant (Pr) mice and in (D) non-reproductive organs for pregnant and non-pregnant (NP) mice. (E, F) Cytokines and chemokines were measured in serum, with red and blue representing high and low levels of expression, respectively. (G, H) For two immunogenic LNPs (503O_8,12_ and 200O_i10_) and two immunoquiescent LNPs (306O_10_ and 306O_i10_), efficacy following IV administration was measured in non-reproductive organs as well as in (I-K) the placenta and fetus. Heatmap colors in panels E and K correspond to median values. Error bars represent s.e.m. n = 3 dams per treatment group, with *, **, ***, and **** representing p < 0.05, 0.01, 0.001, and 0.0001, respectively according to one-way ANOVA with Dunnett’s post-hoc analysis.

As previously shown (21, 36, 37), the route of delivery in non-pregnant mice impacted organ tropism and overall efficacy (Fig. 4D). For example, most luciferase expression for lymphatic tissue, the pancreas, and the liver occurred for IM, IP, and IV delivery, respectively. These route-related differences in organ tropism extended to pregnant mice (Fig. 4D). Interestingly, pregnancy itself impacted organ tropism for IM and IP injections. For example, intramuscular delivery increased lymph node expression by 8-fold in pregnant dams compared to non-pregnant mice, whereas IP delivery reduced it by 4-fold. Pregnancy, however, did not significantly impact the organ-specific protein expression following IV delivery. Because LNPs showed efficacy changes only in lymphoid organs of pregnant mice, we wondered if the shifts were related to the immune response elicited by different administration routes. We therefore measured the blood levels of 11 cytokines and chemokines using Luminex, four hours post-injection. IFNγ and IL-4 were not detected in any sample. Interestingly, IM and IP routes that altered organ-tropism prompted higher cytokine and chemokine production than IV in both pregnant and non-pregnant mice (Fig. 4E). Our results suggest that immunogenic routes can alter protein expression during pregnancy.

To examine whether altered tropism in pregnant vs non-pregnant mice is induced by a differential innate immune response, we asked if immunogenic LNPs cause similar shifts in organ distribution. For this, we compared the immunogenic ionizable lipids 503O_8,12_ and 200O_i10_ and the immunoquiescent lipids 306O_10_ and 306O_i10_. All lipids formed monodisperse LNPs that were 100-120 nm in size (Fig. S2A) and elicited >90% mRNA encapsulation efficiency (Fig. S2B). Further, innate immunogenicity was determined using a Raw-Blue macrophage NF-κB reporter cell line (Fig. S3). We then confirmed that 503O_8,12_ and 200O_i10_ were immunogenic *in vivo* in both pregnant and non-pregnant CD-1 mice following IV injection (Fig. 4F). Additionally, expression of the pro-inflammatory cytokine IL-1β was substantially lower in pregnant dams (Fig. 4F). Interestingly, pregnancy altered organ tropism for immunogenic lipids but not for immunoquiescent lipids (Fig. 4G, 4H). All four lipids induced placental delivery without causing luciferase expression in the fetus (Fig. 4I–K).

Together, these data indicate that LNPs deliver mRNA to the placenta in mice without penetrating the placental barrier, regardless of administration route. Further, the degree of inflammation in the mouse affected organ tropism.

### LNPs transfected placental trophoblasts, endothelial cells, and immune cells

Having shown that LNPs deliver mRNA to the placenta, we next determined which cell layers were transfected. The mouse placenta contains three layers – decidua, junctional zone, and labyrinth (Fig. 5B). The decidua (Dec) is the mucosal lining of the maternal uterus where the embryo implants (38). The labyrinth (Lab) is the primary site of nutrient and gas exchange and is enriched in trophoblasts, endothelial cells, and immune cells (39, 40). The junctional zone (JZ) contains glycogen cells and several trophoblast subtypes, such as trophoblast giant cells and spongiotrophoblasts, and performs endocrine functions, such as hormone regulation, glucose production, and cell metabolism (39, 41). Together, trophoblasts, endothelial cells, and placental immune cells facilitate nutrient and gas exchange, and protect the developing fetus from maternal immune response and invading pathogens.

**Figure 5:**
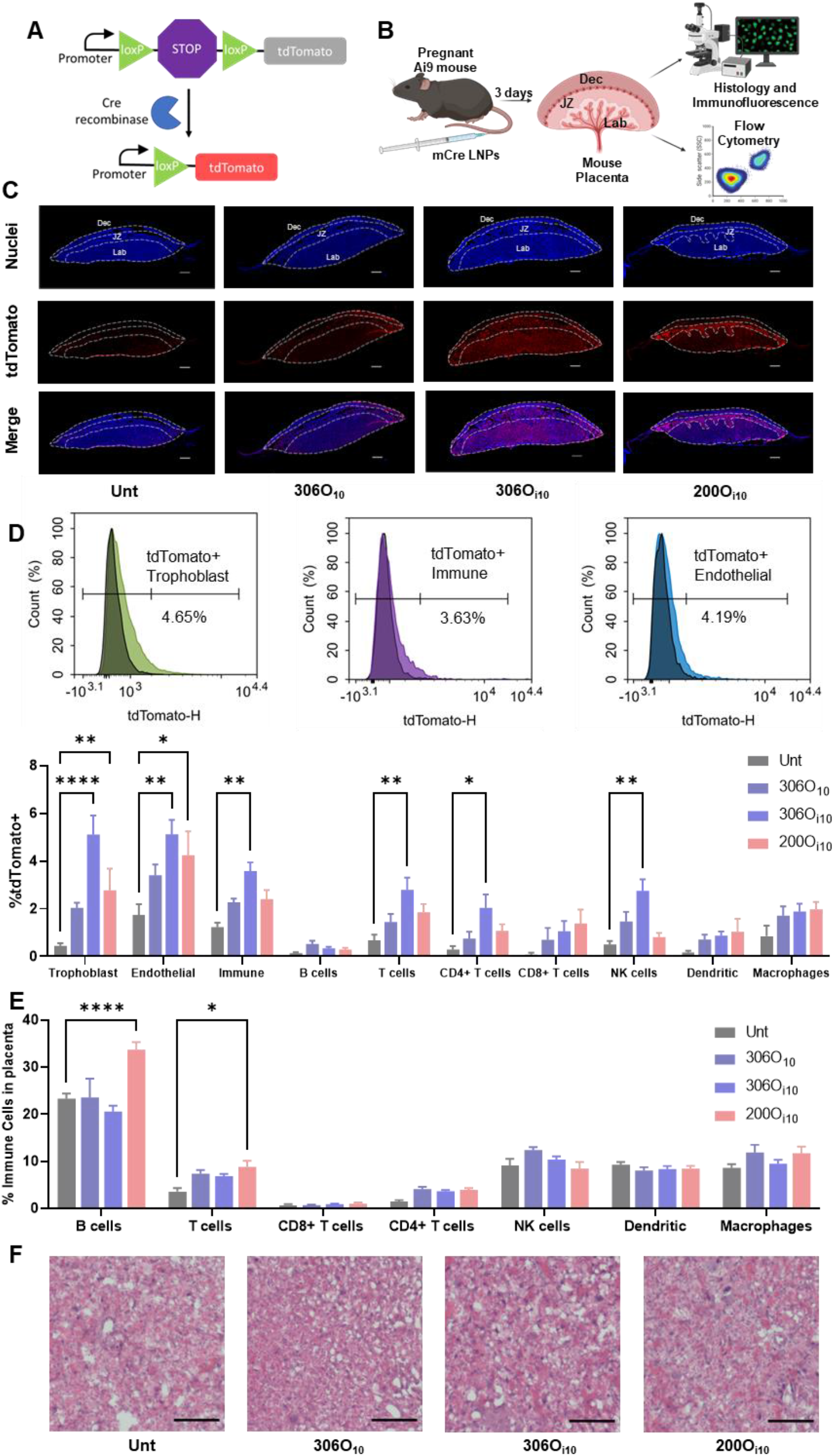
The lipid nanoparticle 306O_i10_ delivers mRNA to placental trophoblasts, endothelial cells, and immune cells in mice. (A) The delivery of mRNA encoding Cre recombinase (mCre) induces Cre-Lox recombination in Ai9 mice and causes tdTomato expression in transfected cells. (B) LNPs formulated with three different ionizable lipids encapsulating mCre were IV-injected in Ai9 mice at 1 mg/kg on the 14^th^ day of pregnancy. After three days, mice were euthanized, and their placentas were analyzed using immunofluorescence microscopy, flow cytometry, and histology. (C) Placentas were stained to distinguish nuclei (blue) and tdTomato (red) and visualized using an immunofluorescent microscope. The decidua (Dec), junctional zone (JZ), and placental labyrinth (Lab) are shown using dotted lines. All scale bars are 0.5 mm. (D) tdTomato expression and (E) immune cell infiltration was measured in isolated placental cells using flow cytometry. (F) Placental morphology was visualized using H&E staining. All scale bars are 50 μm. Error bars represent s.e.m. n = 3 dams per treatment group, with *, **, ***, and **** representing p < 0.05, 0.01, 0.001, and 0.0001, respectively, according to two-way ANOVA with Tukey’s post-hoc analysis compared to untreated controls.

To identify transfected placental cell layers, we used transgenic Ai9 mice that carry a tdTomato gene downstream of a *loxP*-STOP cassette. In naïve Ai9 mice, the cassette prevents tdTomato expression. However, the presence of the recombinase protein, Cre, delivered through LNPs, would induce permanent tdTomato expression in transfected cells by removing the *loxP*-STOP cassette (Fig. 5A). For this study, we IV-injected the two LNPs with the lowest levels of immunogenicity containing Cre recombinase mRNA on the 14^th^ day of pregnancy. Mice were euthanized after three days, and placentae were analyzed for tdTomato expression (Fig. 5B).

First, qualitative assessment using immunofluorescence microscopy revealed that that 200O_i10_ and 306O_i10_ elicited higher tdTomato expression than 306O_10_ (Figs. 5C, S4). Interestingly, the site of protein expression differed for both LNPs. 306O_i10_ showed uniform delivery to the placental labyrinth and junctional zone; however, 200O_i10_ induced higher expression in the junctional zone, and 306O_10_ showed low expression levels in all areas (Fig. 5C). Flow cytometry data confirmed the superior efficacy of 306O_i10_ LNPs for all cell types (Fig. 5D). Delivery via this LNP increased the fraction of tdTomato positive cells 10-, 4-, and 3-fold for trophoblasts, endothelial cells, and immune cells, respectively, compared to untreated mice. Among immune cells, 306O_i10_ primarily transfected CD4+ T cells and NK cells.

To assess LNP safety, we measured immune cell infiltration into placental tissue and evaluated tissue morphology. Interestingly, the 200O_i10_ LNPs that prompted an innate immune response in CD-1 mice (Fig. 4F) also caused higher levels of adaptive immune cells in the placenta. Specifically, 200O_i10_ LNPs caused a 1.5-fold increase in B cells and 2.4-fold increase in T cells compared to untreated animals (Fig. 5E). The immunoquiescent LNPs 306O_10_ and 306O_i10_ (Fig. 5E) did not alter the fraction of immune cells in the placenta. Histological staining showed no gross morphological changes in the placental labyrinth, indicating that LNPs don’t damage the placental vasculature (Fig. 5F).

### 306 LNPs do not affect pup development

Although our data show no mRNA delivery to the fetus, it is possible that placental transfection nonetheless affects pregnancy outcome and maternal and pup health. To assess this, we IV-injected LNPs encoding Cre recombinase in pregnant Ai9 mice and measured outcomes and monitored pup health for three weeks post-delivery. The use of the Cre-expressing LNP also allowed us to track LNP deposition in maternal and fetal tissues.

No dam or pup mortality was noted after LNP administration. Further, all pups were born after a 19-day gestation period, ruling out premature births. Mice were euthanized three weeks after birth to assess LNP delivery. Strikingly, tdTomato expression in the livers of LNP-treated dams was visible to the naked eye (Fig. 6A). Quantification by IVIS showed >300-fold protein expression in the liver for all LNPs compared to untreated controls (Fig. 6B). Further, all maternal organs except for the brain expressed tdTomato. In contrast, the livers of pups born to LNP-treated dams did not visually express tdTomato (Fig. 6C), with IVIS measurements confirming no tdTomato expression in pups as a function of organ (Fig. 6D). This is consistent with our luciferase delivery data in CD-1 mice that showed no expression in fetuses (Fig. 3C, 4C, 4K).

**Figure 6:**
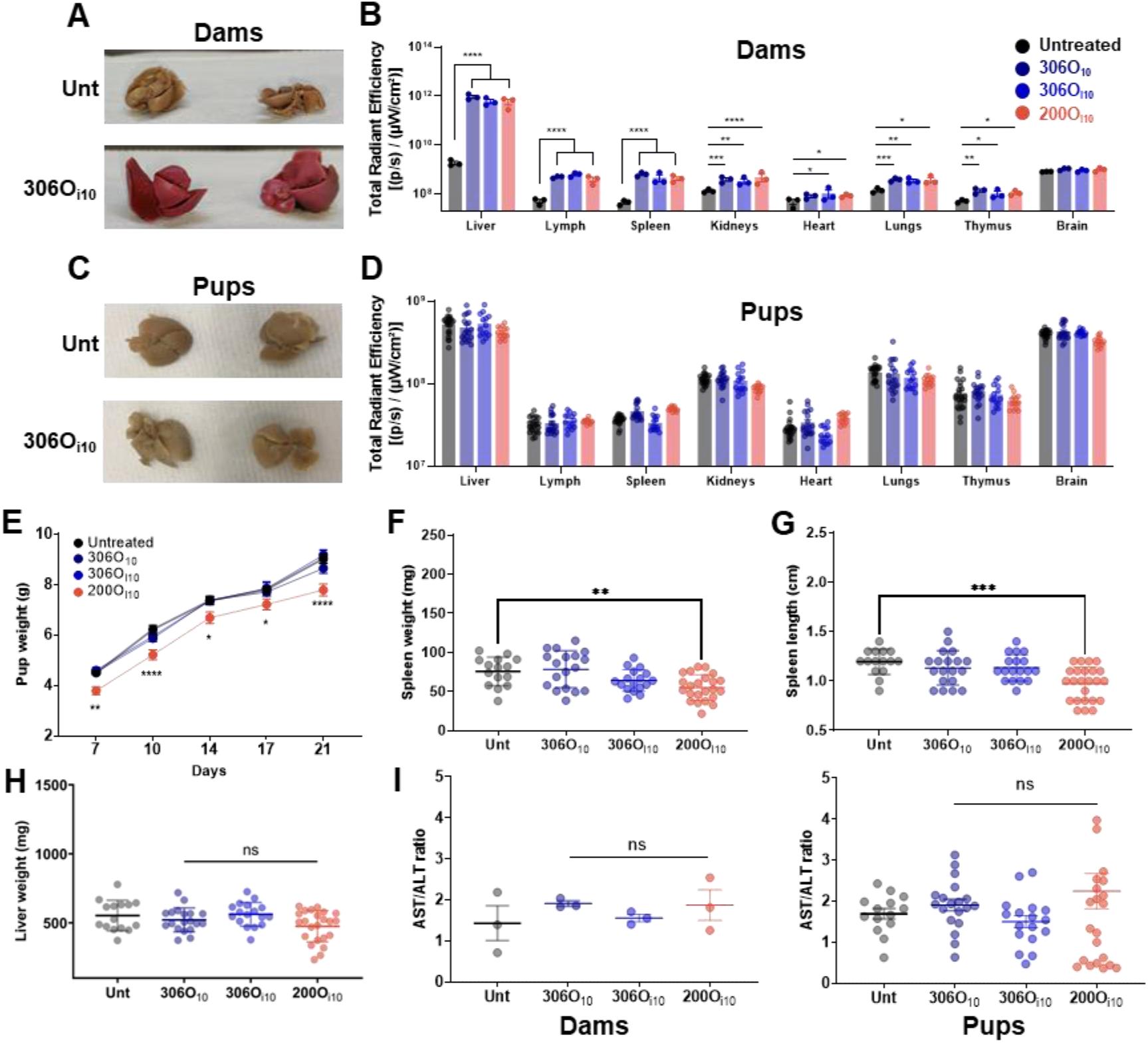
306O_10_ and 306O_i10_ do not affect pup development. LNPs encapsulating Cre recombinase mRNA (mCre) were injected in pregnant Ai9 mice on the 14^th^ day of pregnancy. Three weeks after birth, mice were euthanized, and the organs of (A, B) dams and (C, D) pups were analyzed visually or using IVIS. Pup development was assessed by measuring their (E) body weight over three weeks, and (F) length of their spleen, and the weights of their (G) spleen and (H) liver after euthanasia. (I) AST and ALT were measured in serum of dams and pups to assess hepatotoxicity. Error bars represent s.e.m. n = 3 dams per treatment group, and 3–8 pups per dam with *, **, ***, and **** representing p < 0.05, 0.01, 0.001, and 0.0001, respectively, according to one-way ANOVA with Tukey’s post-hoc analysis compared to untreated controls.

In addition to delivery, we monitored pup health. Interestingly, inflammatory 200O_i10_ LNPs slowed pup growth, as assessed by overall weight and spleen weight and length (Fig. 6E–G). There was no reduction in liver weight (Fig. 6H). Furthermore, the ratio of the liver enzymes AST and ALT were not elevated in either dams or pups (Fig. 6I), indicating no signs of hepatotoxicity.

## Discussion

In this work, we sought to develop safe and potent RNA delivery vehicles that may be used to treat a broad range of pregnancy-related conditions. To this extent, we assessed RNA delivery efficacy, toxicity, placental transport, and immunogenicity of multiple LNP structures and clinically relevant administration routes.

Structure-efficacy screening revealed that the amine headgroup dictates delivery in placental BeWo b30 human trophoblasts (Fig. 1A). Interestingly, the headgroups 306, 500, 503, 514, and 516, which are synthetic analogs of the naturally occurring polyamines spermine and spermidine, showed highest efficacy. In mammalian cells, spermine and spermidine are critical for cell growth, differentiation, and survival (42). They regulate gene expression by binding to DNA and RNA, initiating translation, and interacting with transcription factors (42, 43). Because they complex nucleic acids with high affinity, spermine and spermidine containing lipids have been used in commercial transfection reagents, such as Transfectam® and Lipofectamine® (44), and to deliver DNA (45, 46), siRNA (47, 48), and mRNA (49). Together, these reports and our results suggest that polyamine-derived lipids form potent RNA delivery vehicles. Further, because polyamine synthesis is critical for fetal growth and is upregulated in the placenta (50), we believe that polyamine-derived lipids can effectively deliver mRNA to the placenta.

In contrast to delivery, we found that LNP hemolytic activity was governed by the lipid tail (Fig. 1B). Specifically, lipids containing methylated branches elicited higher hemolysis. We suspect that these branched ionizable lipids disrupt RBC membrane integrity by altering its mechanical properties. Branched lipids are known to disrupt lipid packing by increasing surface area per lipid and elevating membrane microviscosity by restricting lipid motion (51). In the acidic endosome, this disruption is desirable because it facilitates endosomal escape and mRNA delivery to the cytosol; however, membrane disruption in the blood at physiological pH will damage circulating cells, such as red blood cells.

Using viable LNPs from *in vitro* screening, we observed high luciferase delivery in pregnant and non-pregnant CD-1 mice. Interestingly, LNPs elicited route- and structure-dependent shifts in protein expression in the spleen and lymph nodes of pregnant mice (Fig. 4). These therapeutic modalities may require dose adjustment in pregnant people. Further, cytokine and chemokine profiles demonstrated that shifts in organ tropism correlated with immunogenic routes and structures. Particularly, intravenous LNP delivery suppressed IL-1β in pregnant mice. Interleukin family cytokines, including IL-1α and IL-1β are potent mobilizers of innate immune cells, such as monocytes, macrophages, and neutrophils (52). Because patrolling myeloid cells are among the first cells to uptake LNPs *in vivo* (53), we suspect that IL-1β suppresses infiltration of transfected myeloid cells in the spleen and lymph node, and reduces efficacy in these organs.

In addition to tropism, we found that disrupting immune homeostasis deleteriously affects pup development (Fig. 6). Immunogenic 200O_i10_ LNPs elevated B and T cell infiltration in the placenta and impeded pup growth. Although the phenotype of these cells is not known, we suspect that these adaptive immune cells induce inflammatory responses which may adversely affect fetal development. Previously, inflammatory cytokines produced by infiltrating T cells have been shown to affect pup development (54).

Our results highlight that immune homeostasis is critical for fetal safety. Because 306O_i10_ LNPs are safe during pregnancy, they can be employed for a myriad of prophylactic and therapeutic applications. For example, prophylactic mRNA vaccines encoding targeting lethal pathogens, such as influenza, can be administered intramuscularly during pregnancy and protect fetuses from pathogen-induced congenital anomalies (55). Immune evasive viruses, such as HIV, which elicit neutralizing antibodies in a rare fraction of infected individuals (56), might benefit from passive vaccination strategies. Here, mRNA-LNPs encoding broadly-neutralizing antibodies, such as VRC01 (57, 58), BG18 (59), PGT121 (60), can be translated by hepatocytes after IV delivery, and neutralize HIV strains. Since antibodies can cross the placenta via FcRn-mediated transcytosis (61), vertically-transferred maternal antibodies can protect the fetus from adverse outcomes of perinatal HIV transmission (62).

For treating maternal disorders where placental delivery is relevant, intravenous mRNA delivery may be the preferred route. We found that 306O_i10_ delivered mRNA to the placental junctional and labyrinthine zone and transfected several placental cell types (Fig. 5). Therefore, its therapeutic applications can extend to disorders that involve multiple cell types, such as preeclampsia. For example, protein supplementation by delivering mRNA encoding angiogenic proteins, such as vascular endothelial growth factor or placental growth factor, can promote placental vascularization that is perturbed during preeclampsia (63, 64). Further, silencing anti-angiogenic proteins, such as soluble fms-like tyrosine kinase-1 (sFlt-1) that are overexpressed by trophoblasts during preeclampsia is an attractive therapeutic strategy (2). Here, transient gene knockdown of sFlt-1 using CRISPRi based approaches can suppress sFlt-1 levels in the placenta and bloodstream and ameliorate preeclampsia (65, 66).

In this study, we identified potent LNP structures that deliver mRNA to both the placenta and non-reproductive maternal organs without fetal delivery. To our knowledge, this is the first study that provides a mechanistic basis for pharmacodynamic shifts in LNP delivery during pregnancy. Further, we demonstrate that LNP immunogenicity is a critical parameter for designing safe therapeutics for pregnancy. Together, we anticipate that this work will propel the design of potent LNPs that will establish new pregnancy-safe treatment paradigms.

## Methods

### Materials

Amines were purchased from Sigma Aldrich (St. Louis, MO), CHESS Organics (Mannheim, Germany), Acros Organics (Fair Lawn, NJ), and Alfa Aesar (Haverhill, MA). Acrylates were sourced from Scientific Polymer Products (Ontario, NY), Hampford Research, Inc. (Stratford, CT), Monomer-Polymer and Dajac Labs (Trevose, PA), and Sartomer (Warrington, PA). 1,2-dioleoyl-sn-glycero-3-phosphoethanolamine (DOPE) and PEG-DMPE were obtained from Avanti Polar Lipids (Alabaster, AL). Cholesterol and sodium citrate were sourced from Sigma Aldrich (St. Louis, MO). 3500 MWCO dialysis cassettes were purchased from Thermo Fisher (Waltham, MA). mRNA encoding Cre recombinase and firefly luciferase were purchased from TriLink BioTechnologies (San Diego, CA). D-Luciferin was obtained from PerkinElmer (Waltham, MA). BeWo cells were obtained from ATCC, and BeWo b30 cells were a generous gift from the laboratory of Dr. Alan Schwartz (Washington University, St. Louis, MO). DMEM/F12, fetal bovine serum (FBS), donkey serum, Penicillin/Streptomycin, phosphate buffered saline (PBS), Hanks’ Balanced Salt Solution (HBSS), and HEPES were sourced from Thermo Fisher (Waltham, MA). Lipopolysaccharide from *E. coli* 0111:B4 (LPS) was purchased from Sigma-Aldrich (St. Louis, MO). Raw Blue™ cells, Normocin™, Zeocin™, and the QUANTI-Blue™ assay kit were obtained from Invivogen (San Diego, CA). BrightGLO™ was purchased from Promega (Madison, WI). Human Red blood cells were sourced from Innovative Research (Novi, MI), and Triton-X 100 was obtained from Alfa Aesar (Haverhill, MA). Collagenase Type I was purchased from Worthington Biochemical Corporation (Lakewood, NJ), and DNAse I was purchased from Thermo Fisher (Waltham, MA). Formaldehyde was sourced from VWR (Radnor, PA). Hoescht, AF488 phalloidin, AF594 anti-ZO-1, goat anti-RFP, and AF647 donkey anti-goat IgG (H+L) were purchased from Thermo Fisher (Waltham, MA). Mouse Luminex Discovery Assay kit configured for CCL2, CXCL1, CXCL2, IFN-γ, IL-1β, IL-2, IL-4, IL-6, IL-10, TNFα, and VEGF was purchased from R & D Systems (Minneapolis, MN). 10X RBC lysis buffer and antibodies used for flow cytometry were sourced from Biolegend (San Diego, CA). Hematoxylin, Eosin, Scott’s Tap Water Substitute, and xylene were sourced from Millipore Sigma (Burlington, MA). 2,4-Dinitrophenylhydrazine was purchased from Fisher Scientific (Waltham, MA). Sodium hydroxide, L-alanine, L-aspartic acid, α-Ketoglutaric acid, and sodium pyruvate were obtained from Millipore Sigma (St. Louis, MO).

### Nanoparticle formulation and characterization

Lipids were synthesized using Michael addition chemistry by combinatorically reacting an amine with an alkyl acrylate at 90 °C for three days (67). Nanoparticles were formulated as described previously (67). Briefly, the ionizable lipid, DOPE, cholesterol, and C14-PEG2000 were dissolved in ethanol (90% v/v) and 10 mM sodium citrate (10% v/v) at a molar ratio of 35:16:46.5:2.5. The mRNA was dissolved in 10 mM sodium citrate. Nanoparticles were formed by rapidly pipetting an equal volume of the lipid solution into the RNA solution. Lipids were added at a 10:1 total lipid:mRNA w/w ratio. All nanoparticles injected into mice were dialyzed in PBS for 90 min using 3500 MWCO dialysis cassettes. Nanoparticle size and dispersity were assessed using a Malvern Zetasizer Nano ZSP (Malvern, UK). Further, mRNA entrapment was measured using the Quant-iT RiboGreen assay (Thermo Fisher).

### *In vitro* mRNA delivery

BeWo or BeWo b30 cells were cultured in DMEM/F12 supplemented with 10% fetal bovine serum and 1% penicillin/streptomycin and incubated at 37 °C and 5% CO_2_. Cells were seeded at 100,000 cells/well in white flat bottom 96-well plates (Greiner Bio-One, Monroe, NC). After 48 hrs, LNPs formulated with firefly luciferase mRNA were added to cells at a dose of 100 ng mRNA per well. Twenty-four hrs later, luciferase activity was measured using a Bight-Glo™ Luciferase Assay System (Promega, Madison, WI) as per the manufacturer’s protocol.

### Hemolysis

Human Red blood cells (Innovative Research, Novi, MI) were diluted in PBS to make a 4% v/v RBC solution. 100 μL RBC solution was seeded in clear round bottom 96-well plates (Corning, Corning, NY), and an equal volume of blank LNPs (i.e., containing no mRNA) formulated at 0.08 mg/mL equivalent mRNA concentration (2500 ng mRNA/well) was added to the solution. PBS and 1% Triton-X were used as negative and positive controls, respectively. The solution was incubated for 90 min at 37 °C. Cells were centrifuged at 500 RCF, and 100 μL supernatant was transferred to a transparent flat bottom 96-well plate (Corning, Corning, NY). Absorbance was measured at 540/640 nm using a Synergy H1 microplate reader (BioTek Instruments, Winooski, VT).

### *In vitro* permeability

BeWo or BeWo b30 cells were seeded on 0.4 μm polycarbonate Transwell® permeable supports (Corning, Corning, NY) at a density of 100,000 cells/well. Media was changed daily, and monolayer formation was monitored by measuring *trans-* epithelial electrical resistance (TEER) using a Millicell voltohmmeter (MilliporeSigma, Burlington, MA). TEER values above 150 Ω-cm^2^ indicated confluent monolayer formation. On the day of the experiment, Transwell^®^ inserts were transferred into transport buffer (HBSS with 12.5 mM glucose and 25 mM HEPES) and equilibrated at 37 °C for one hour. To assess transport, Cy5-labeled mRNA or fluorescent marker molecules were added to the apical side of the inserts. Groups included Cy5-labeled mRNA (naked and LNP-formulated), fluorescein, and 4 kDa and 10 kDa FITC-dextrans at final concentrations of 2.5 μg/mL, 25 μM, and 100 μg/mL, respectively. Transwells® were then incubated at 37 °C and 25 RPM for 2 hrs in a New Brunswich™ Innova® 40 shaker (Eppendorf, Enfield, CT). Samples were collected every hour from the basolateral side. Cy-5 and fluorescein/FITC fluorescence was measured at 650/670 nm and 485/530 nm, respectively, using a plate reader. Apparent permeability (P_app_) was calculated using the equation:

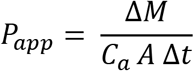

where P_app_ is the apparent permeability, ΔM is the mass of the fluorescent molecule transported into the basolateral compartment, C_a_ is the concentration of the fluorescent molecule on the apical side, A is the area of the insert, and Δt is the time interval.

### Immunocytochemistry

BeWo b30 monolayers were rinsed with PBS fixed in 3.7% formaldehyde. Samples were permeabilized with 0.2% Triton-X and blocked with 5% FBS in PBS. Monolayers were stained with 1 μg/mL Hoescht, 165 nM AF488 phalloidin, and 1 μg/mL AF594 anti-ZO-1 antibody for at least an hour. Monolayers were mounted on slides and imaged using a fluorescence microscope.

### *In vivo* mLuc delivery

All mouse experiments were approved by the Institutional Animal Care and Use Committee (IACUC) at Carnegie Mellon University under protocol number PROTO201600048. Pregnant CD-1 mice were purchased from Charles Rivers and lipid nanoparticles encapsulating luciferase mRNA were administered on the 14^th^ day of pregnancy. Four hours after LNP injection, mice were intraperitoneally injected with 130 μL D-Luciferin (30 mg/mL in PBS). Fifteen minutes later, mice were euthanized using CO_2_ asphyxiation, and organs and blood were collected. Organs were imaged to measure luminescence using IVIS® (PerkinElmer, Waltham, MA). Total luminescent flux (photons/second) was determined using region of interest analysis using the Living Image® software. Blood samples were centrifuged at 12,000 RPM for 10 min, and cytokines and chemokine levels were measured in serum using Luminex as per the manufacturer’s instructions.

### *In vivo* mCre delivery

Pregnant Ai9 mice were obtained from an institutionally managed animal colony. Mice were dosed with LNPs encapsulating mRNA encoding Cre Recombinase on the 14^th^ day of pregnancy. For placenta delivery experiments, mice were IV-injected with 1 mg/kg LNPs. After three days, mice were euthanized, and placentae were either processed for flow cytometry or immediately fixed in 3.7% formaldehyde for histological and immunofluorescence analysis. For pup growth experiments, pregnant mice were IV-injected with 0.5 mg/kg LNPs. Twenty-one days after birth, dams and pups were euthanized, and their organs and blood were collected. Organs were imaged using IVIS® and Total Radiant Efficiency (p/s)/(μW/cm^2^) was calculated using region of interest analysis using the Living Image® software. Blood was centrifuged, and serum was used to measure aspartate transaminase and alanine transaminase activity.

### Flow cytometry

Placentas were harvested and placed in DMEM at 4 °C after removing the decidua. Placentas were digested for 35 min in 1 mL digestion buffer (DMEM, 500 units/mL collagenase I, and 8 units/μL DNAse I). Cells were passed through a 70 μm cell strainer and washed with cold DMEM. Cells were centrifuged at 400 g, and Red blood cells were lysed using 1X RBC lysis buffer (Biolegend). Cells were stained with fixable yellow viability dye (1:1000 dilution) for 15 min at 4 °C as per the manufacturer’s instructions. After staining, cells were fixed with 3.7% formaldehyde for 10 min and blocked with 5% FBS. Cells were stained with an antibody panel for lymphoid cells, myeloid cells, or trophoblasts. In the lymphoid panel, cells were stained with Fc block, PerCP anti-mouse CD45 (clone: 30-F11), PE-Cy7 anti-mouse CD4 (clone: GK-1.5), BV605 anti-mouse CB8a (clone: 53-6.7), BV421 anti-mouse CD19 (clone: 6D5), BV785 anti-mouse CD3 (clone: 17A2), and APC anti-mouse CD49b (clone: DX5). In the myeloid panel, cells were stained with Fc block, PerCP anti-mouse CD45 (clone: 30-F11), APC anti-mouse CD11c (clone: N418), BV650 anti-mouse CD11b (clone: M1/70), and BV785 anti-mouse F4/80 (clone: BM8). In the trophoblast panel, cells were stained with Fc block, PerCP anti-mouse CD45 (clone: 30-F11), BV785 anti-mouse CD31 (clone: 390), and polyclonal rabbit anti-cytokeratin 7 followed by goat anti-rabbit AF647 secondary. All antibodies were added at a concentration of 2 μg/mL for 35 min. After staining, cells were resuspended in 5% FBS and analyzed by flow cytometry using a NovoCyte 3000 (ACEA Biosciences, San Diego, CA). Flow cytometry data were analyzed using NovoExpress software.

### Immunofluorescence and Histology

Placentas were dissected and immediately fixed in 3.7% formaldehyde for 2 days at 4 °C. Placentas were washed with PBS and dehydrated by incubating in 15% and 30% sucrose for 12 hrs each. Samples were embedded in optimal cutting temperature compound (Tissue Tek, Torrance, CA), snap frozen using liquid nitrogen, and sliced into 10 μm sections on ColorFrost plus microscope slides (Fisher Scientific, Waltham, MA). For immunofluorescence imaging, samples and subject to antigen retrieval by incubating in 10 mM citrate buffer / 0.05% Tween-20 (pH 6) at 95 °C followed by autofluorescence reduction in 10 mM copper sulfate / 50 mM ammonium acetate at 37 °C for 20 min each. Samples were blocked and stained with 5 μg/mL anti-RFP for 12 hrs, washed 3X with PBS, followed by staining with 5 μg/mL AF647 donkey anti-goat IgG (H+L) for 1 hr. Samples were counterstained with 100 ng/mL DAPI for 15 min, mounted on cover slides, and imaged using a fluorescent microscope. For histology, samples were stained with hematoxylin for 4 min, rinsed with 0.3% HCl in 70% EtOH, rinsed in Scott’s tap water, and stained with eosin for 2 min. Slides were washed with PBS between each step. After staining, slides were dehydrated in 95% and 100% EtOH, cleared with xylene, mounted on cover slides, and imaged using a brightfield microscope.

### Alanine aminotransferase (ALT) and Aspartate aminotransferase (AST) activity

Serum ALT and AST activities were measured using the Reitman-Frenkel method(68). Briefly, samples were diluted 1:1 with PBS, and 10 μL diluted samples were incubated with 50 μL solution containing 2mM α-Ketoglutaric acid and 200 mM amino acid at 37 °C for 60 min. L-alanine and L-aspartic acid were used as the amino acids for ALT and AST activity, respectively. Sodium Pyruvate was used to generate a standard curve. After an hour, 50 μL 1 mM 2,4-dinitrophenylhydrazine in 1 M hydrochloric acid was added and incubated at 37 °C for 30 minutes. The reaction was stopped by adding 200 μL 4N NaOH, and absorbance was measured at 510/660 nm using a plate reader.

