## Supplemental Information for "Lipid nanoparticle structure and delivery route during pregnancy dictates mRNA potency, immunogenicity, and health in the mother and offspring"

### Supplementary Information

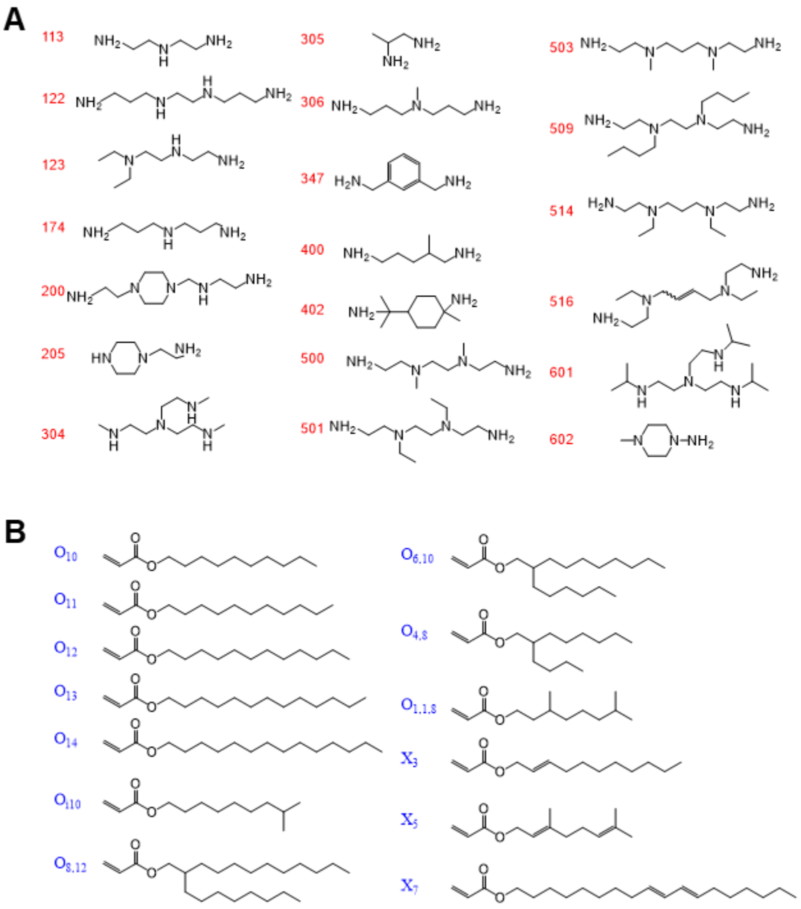


**Figure S1:** A library of 260 ionizable lipids was synthesized by combinatorically reacting (A) 20 amine heads and (B) 13 acrylate tails via Michael addition chemistry.

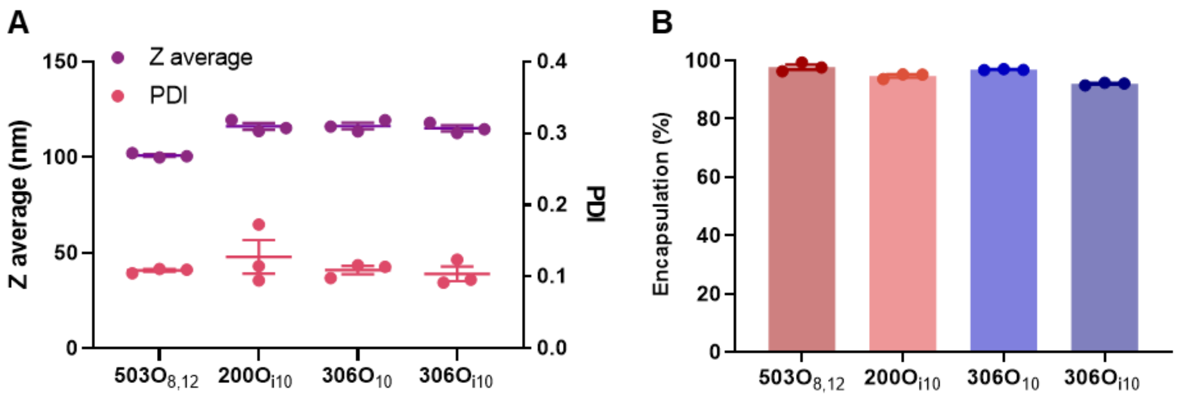


**Figure S2: Lipids form monodisperse nanoparticles, and encapsulate RNA with high efficiency**. LNPs were formulated using four ionizable lipids. (A) Size and dispersity were measured using dynamic light scattering, (B) and mRNA encapsulation efficiency was assessed using Quant-IT RiboGreen assay.

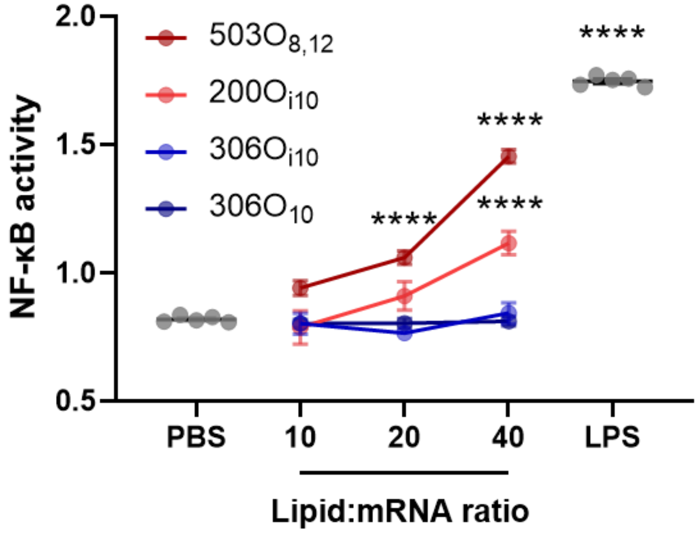


**Figure S3: Two of the four LNPs examined in this study prompted NF-κB activity in Raw Blue™ cells.** Raw Blue™ cells were incubated with LNPs formulated with four distinct ionizable lipids at increasing lipid:mRNA doses, and NF-κB activity was measured using the Quanti Blue™ assay. Error bars represent s.e.m. with $n = 5$ and **** representing $p < 0.0001$, according to one-way ANOVA with Dunnett's post-hoc analysis.

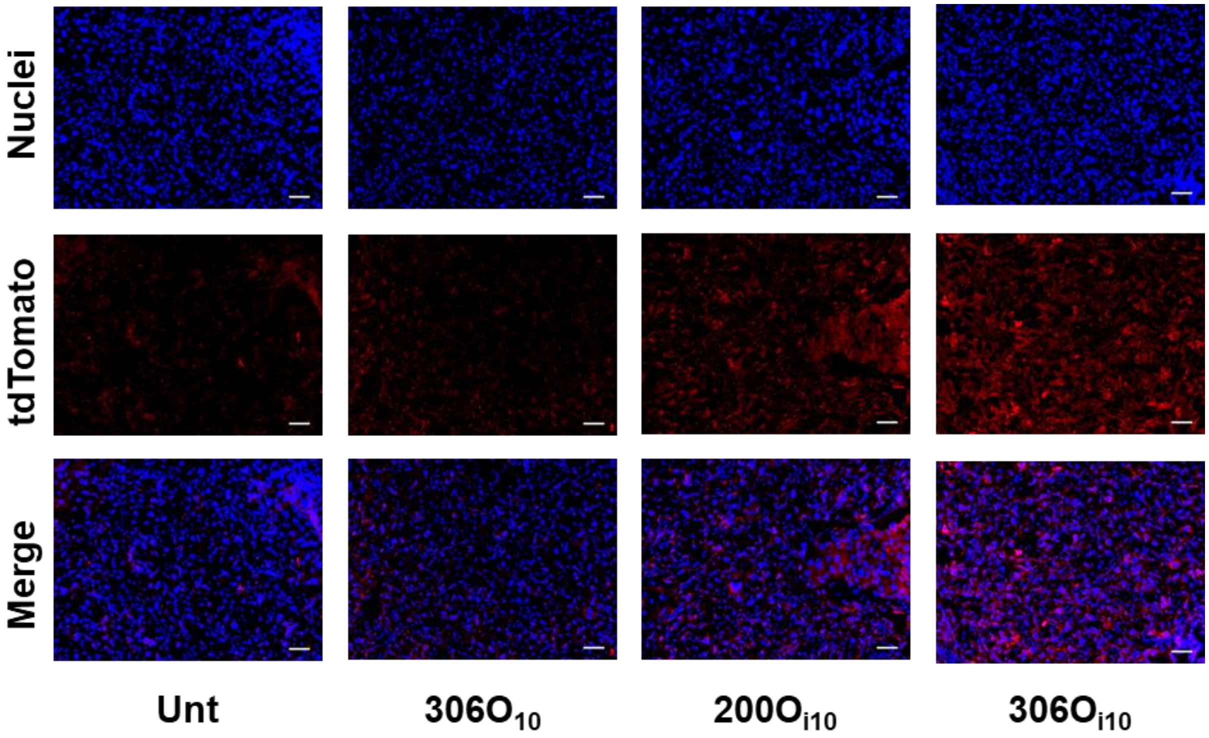


**Figure S4: LNPs deliver mRNA to the placental labyrinth in Ai9 mice.** LNPs encapsulating Cre recombinase (mCre) were injected in pregnant Ai9 mice on the 14th day of pregnancy. After three days, mice were euthanized. (A) Placentas were stained with DAPI (nuclei), and goat anti-RFP and donkey anti-goat AF647 (tdTomato), and placentas were visualized using an immunofluorescent microscope. All scale bars are 20 µm. These are the same data as Fig 4A but at higher magnification.

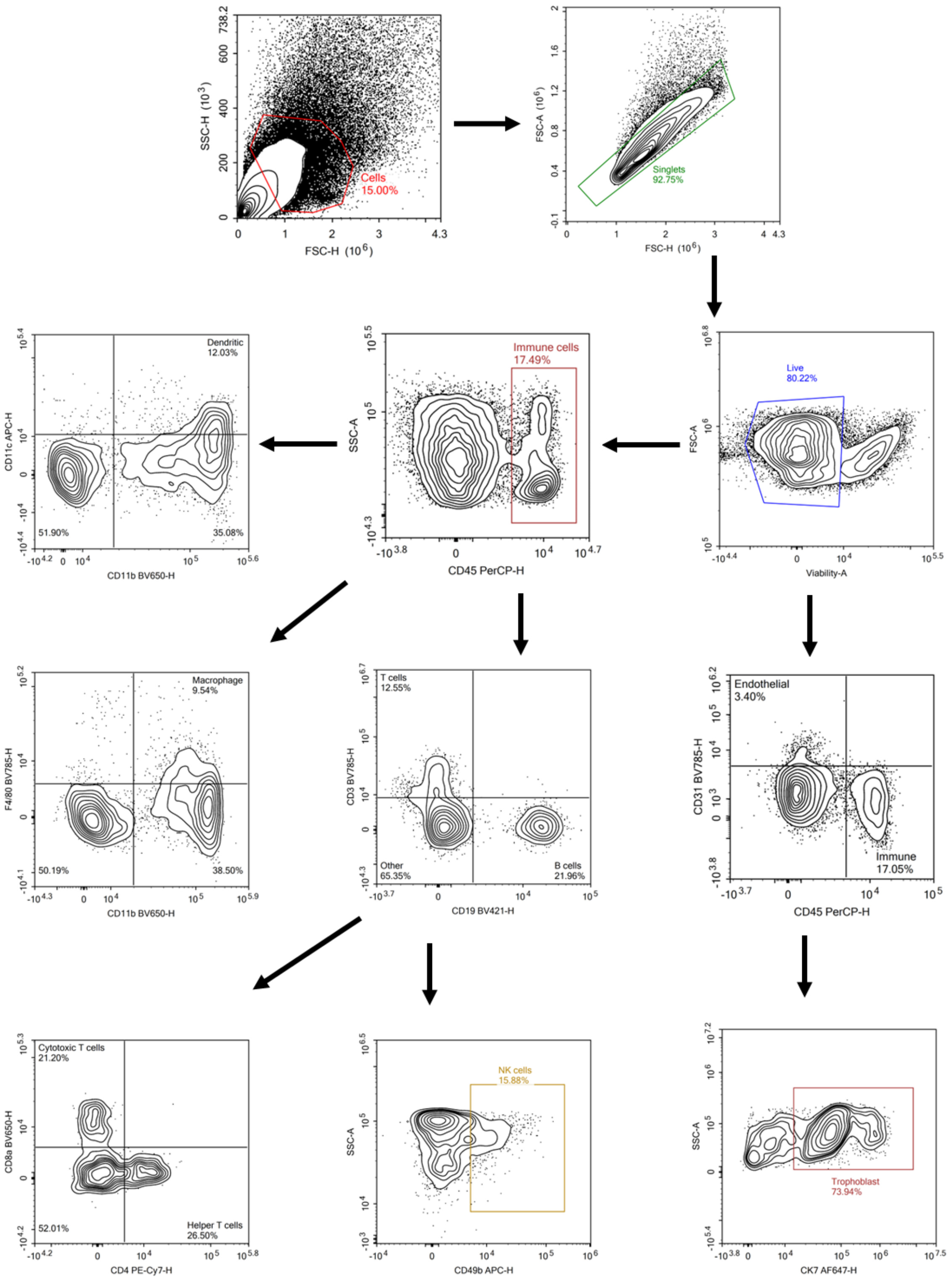


**Figure S5: Gating scheme for flow cytometry.** Placenta were digested and analyzed using flow cytometry. Percentages of trophoblasts (CD45- CD31- CK7+), endothelial cells (CD45- CD31+), immune cells (CD45+ CD31-), B cells (CD45+ CD3- CD19+), T cells (CD45+ CD3+ CD19-), CD4+ T cells, CD8+ T cells, NK cells (CD45+ CD3- CD19- CD49b+) macrophages (CD45+ CD11b+ F4/80+), and dendritic cells (CD45+ CD11b+ CD11c+) were calculated.
